# Pleiotropy in *FOXC1*-attributable phenotypes involves altered ciliation and cilia-dependent signaling

**DOI:** 10.1101/2020.08.13.249334

**Authors:** Serhiy Havrylov, Paul Chrystal, Suey van Baarle, Curtis R French, Ian M MacDonald, Jagannadha R Avasarala, R Curtis Rogers, Fred B Berry, Tsutomu Kume, Andrew J Waskiewicz, Ordan J Lehmann

## Abstract

Alterations to cilia are responsible for a wide range of severe disease; however, understanding of the transcriptional control of ciliogenesis remains incomplete. We evaluated whether ciliary dysfunction contributed to the pleiotropic phenotypes caused by the *Forkhead* transcription factor *FOXC1*. Here, we show that patients with *FOXC1*-attributable Axenfeld-Rieger Syndrome (ARS) have a prevalence of ciliopathy-associated phenotypes comparable to syndromic ciliopathies. We demonstrate that altering the level of Foxc1, via shRNA mediated inhibition and mRNA overexpression, modifies cilia length *in vitro*. These structural changes were associated with substantially perturbed cilia-dependent signaling [Hedgehog (Hh) and PDGFRα] and the altered ciliary compartmentalization of a major Hh pathway transcription factor, Gli2. Analyses of two *Foxc1* murine mutant strains demonstrated altered axonemal length in the choroid plexus with the increased expression of an essential regulator of multi-ciliation, *Foxj1*. The novel complexity revealed in ciliation of the choroid plexus indicates a partitioning of function between these *Forkhead* transcription factors. Collectively, these results support a contribution from ciliary dysfunction to some *FOXC1*-induced phenotypes.

## Introduction

The primary cilium is a sensory organelle, present on most cells, that has essential roles in development and homeostasis. Mutation in genes encoding ciliary proteins, result in an extensive spectrum of phenotypes, in which severe congenital anomalies are over-represented. Consequently, such ciliopathies have been intensively investigated, both uncovering novel pathogenic mechanisms, and providing broader insight into the heritability of disease. For instance, ciliopathy inheritance patterns that were discordant with classical Mendelian models revealed the importance of tri-allelic inheritance (1) and mutational load (2), that in turn represent powerful paradigms for the causality of complex disease.

The importance of cilia stems from their role as a molecular nexus for signal transduction, with primary cilia mediating multiple pathways, including Hedgehog (Hh), and to a variable degree PDGFRα, TGF-β, Hippo and Wnt signalling. Since cilia are indispensable to vertebrate Hh signaling, phenotypes indicative of Hh impairment represent clues that a disorder may be caused by ciliary dysfunction. Examples of such Hh-attributable developmental anomalies include alterations to digit number (syndactyly and polydactyly), the facial skeleton and the midline cerebellum (3, 4). Hydrocephalus, congenital heart disease and renal cysts exemplify additional cilia-associated phenotypes, that reflect roles in mechanosensation, chemosensation, and extracellular fluid movement (5, 6). However, clinical recognition of a ciliopathy can be obscured, by variable phenotypic severity and involvement of individual tissues. An additional complexity is provided by alterations to cilia positioning, that reflect perturbation of cellular orientation across tissue planes, or Planar Cell Polarity (7).

Consistent with the premise that the mechanistic basis of a proportion of ciliopathies may be unrecognized, the majority identified to date are caused by mutations of major effect that induce severe and pediatric-onset disease. Milder alterations to cilia-mediated signaling would be expected to contribute to late-onset phenotypes; however, these remain largely unidentified. A second comparatively undefined area is the regulation of ciliogenesis. Only small numbers of transcription factors have been identified in vertebrates: including the Rfx gene family (*Rfx1-4*) (8–10), and two <u>F</u>orkhead B<u>ox</u> (FOX) family members (*Foxj1* and *Foxn4*) (11–15) that directly regulate development of motile cilia. *FOX* genes have fundamental physiological functions, that extend from angiogenesis and organ development, to cell cycle control (16). Consequently, *FOX* gene mutations induce a diverse disease spectrum, that includes: autism, malignancy, immune deficiency, diabetes, stroke, and speech and language impairment (17–26).

Our study concerns one intensively-studied *FOX* gene, *FOXC1*, which was originally identified as a regulator of organ development (27). Subsequent studies revealed essential roles in arterial specification (28), angio- and somitogenesis (29–31), stem cell quiescence (32, 33) and hematopoietic progenitor formation (34). Heterozygous *FOXC1* mutation and copy number variation (segmental deletion and duplication) cause up to 50% of cases of Axenfeld-Rieger Syndrome (ARS), a pediatric glaucoma-associated disorder whose variable systemic phenotypes include mid-facial hypoplasia, dental anomalies, congenital heart disease and auditory impairment (35–39). In part guided by findings from the murine *Foxc1* mutant, *congenital hydrocephalus* (27, 40), and zebrafish morphants (41), the disease spectrum has extended to include: cerebellar malformations, hydrocephalus, corneal angiogenesis, cerebrovascular disease, and multiple malignancies (25, 42–46). Involvement in stroke and the most severe breast cancer subtype, demonstrates a contribution to common late-onset diseases.

Intrigued that mutation, increased and decreased dosage of a single transcription factor could cause such pleiotropy, and recognizing that some phenotypes were indicative of a ciliopathy, we assessed the hypothesis that *FOXC1* influenced ciliary function. Through evaluation of axonemal length and cilia-mediated signalling *in vitro*, together with studies in two murine mutants, these analyses yielded evidence that Foxc1 modulates ciliation. Developmentally, our findings extend the complexity of the regulation of multi-ciliation in the choroid plexus, and from a clinical perspective, offer an explanation for *FOXC1*’s pleiotropy.

## Results

### Patient phenotypes induced by FOXC1 mutation or copy number variation

We first asked if there was clinical evidence that alterations to *FOXC1* impaired cilia function. Patients with *FOXC1*-attributable Axenfeld-Rieger Syndrome [mutation (n=19), copy number variation (n=22)] (25, 43, 47, 48) were evaluated for traits characteristic of altered cilia-mediated signaling. We established that the prevalence of several ciliopathy-associated phenotypes was substantially elevated as illustrated by the rates of midfacial hypoplasia (27%), congenital heart disease (27%), auditory impairment (24%), cerebellar hypoplasia (15%), or hydrocephalus (8%) (Supplemental Table 1; Figure 1). Multiple phenotypes occurred at or above the rates reported in ciliopathies (ventriculomegaly and hydrocephalus 19%, Joubert Syndrome 23%; congenital heart disease 27%, Nephronophthisis, Bardet-Biedl and McKusick-Kaufman Syndromes 5-18%), while others were less prevalent (cerebellar hypoplasia 15%, Joubert Syndrome 100%) (49–53). This may, in part, reflect the extreme heterogeneity of ciliopathies, as the prevalence of cerebellar hypoplasia (14%) in Bardet-Biedl Syndrome demonstrates (54). A second factor may be that most syndromic ciliopathies are autosomal recessively inherited, with some cases of tri-allelic inheritance (1), while ARS requires a single mutant allele. Amongst the observed phenotypes, polydactyly (Figure 1A) is pathognomonic of altered Hh signaling, which specifies digit number in the developing limb bud (55, 56) where *Foxc1* is expressed (27). Equally, cerebellar anomalies reflect perturbation of the Hh signaling that is essential for cerebellar progenitor cell proliferation (midline cerebellar hypoplasia; Figure 1B). Collectively, the clinical data demonstrate that *FOXC1* mutation or dosage alteration induces variable multi-organ phenotypes, which coincide with the spectrum seen in cilia dysfunction.

**Figure 1.**
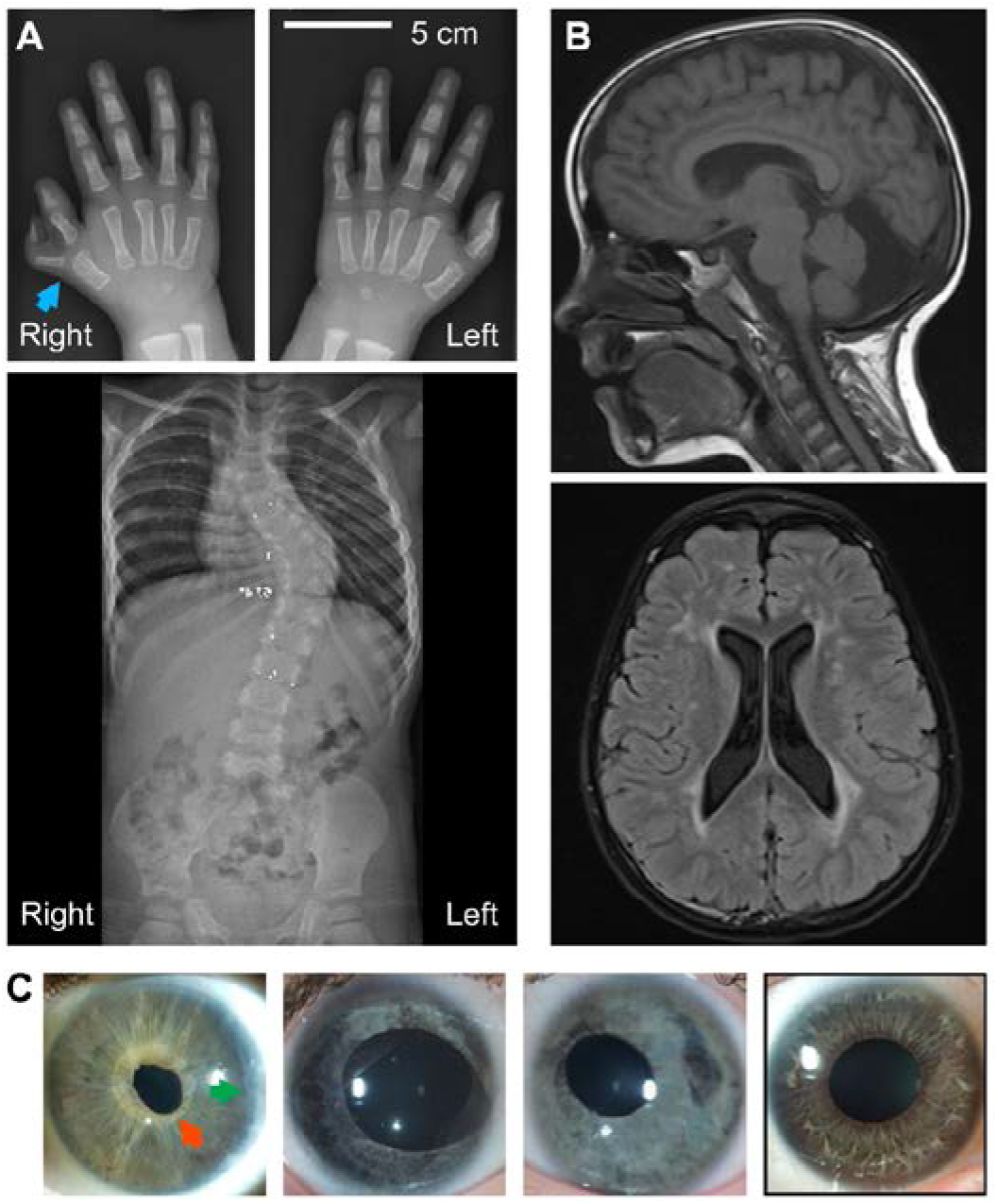
Pleiotropic and variable phenotypes are consistent with ciliary dysfunction. (**A**) Skeletal phenotypes present in the cohort of patients with FOXC1 mutation or copy number variation include scoliosis and pre-axial polydactyly (arrowhead); the duplicated second phalanx of the left thumb was previously surgically removed. The mutation present in this individual (p.D117Tfs64) is predicted to result in loss of two thirds of the FOXC1 protein;. (**B**) CNS phenotypes include cerebellar vermis hypoplasia, posterior fossa enlargement, and lateral ventricular dilation. (**C**) Ocular anomalies include irregular iris sphincter width (red arrow), posterior embryotoxon extending through 3 clock hours (green arrow), and asymmetric irides. Normal iris anatomy for comparison (black box).

### Foxc1 expression influences length of primary cilia in fibroblast, renal and chondrogenic cell lines

We next employed three mammalian cell lines, to determine whether manipulation of the level of Foxc1 impacted cilia structure. Since ARS is frequently caused by segmental duplication and deletion that increase and decrease the copy number of *FOXC1*, the effects of Foxc1 overexpression, and inhibition, were evaluated *in vitro*. In murine fibroblasts (NIH3T3), knockdown of Foxc1 by four independent shRNAs induced mild cilia shortening relative to cells expressing either a non-targeting shRNA or a control plasmid, while, comparable cilia lengthening was observed with stable overexpression of Foxc1 (Figure 2A). To better evaluate the magnitude of these effects, automated image analysis methodology was developed, and validated by comparing manual and automated cilia length measurements in cells treated with a Foxc1-targeting shRNA or vector control (pLKO.1). The mean cilia lengths quantified by the two approaches, were concordant (automated: Foxc1 shRNA 0.85, control 1.0, *P* = 4.5 × 10^−4^; manual: Foxc1 shRNA 0.87, *P* = 1.6 × 10^−3^; Supplemental Figure 1). Measurement of a larger number of cilia revealed that Foxc1 inhibition is accompanied by an altered cilia length distribution in the cell population (*P*_*KW*_ = 7 × 10^−3^; n=7,431 cilia; Figure 2B), with an increased proportion of short cilia (see Methods: shRNA 34 - 44%; controls 24 - 27%; *P* = 1.7 × 10^−4^; Figure 2C), and overall, a 13% shortening relative to controls (6 – 19% for individual shRNAs; Figure 2C). Increased Foxc1 expression induced a 9% lengthening, primarily due to an increased subpopulation of cells with longer cilia (Foxc1 overexpression 58%; controls 50%; *P* = 0.02; n=9,590 cilia; Figure 2C). These data illustrate that automated analysis in large numbers of cells readily resolves mild alterations in cilia length. Quantification of cilia length in a second murine cell line that expresses Foxc1 at a higher level [inner medullary collecting duct cells (IMCD3)], revealed analogous alterations (Foxc1 shRNA inhibition: 30 - 31% shortening, *P* = 7.6 × 10^−7^; overexpression: 10% lengthening, *P* = 6.5 × 10^−5^, Figure 2D-F). Consistent with the reciprocal effects of increased and decreased Foxc1 expression in NIH3T3 and IMCD3 cells, CRISPR/Cas9 mutagenesis of *Foxc1* in a chondrogenic cell line (ATDC5) induced an 11% reduction in cilia length compared to CRISPR-treated control cells (*P* = 3 × 10^−5^; Figure 2G-I). This demonstrates that a targeted loss of function *Foxc1* mutation, recapitulates the effect of shRNA inhibition.

**Figure 2.**
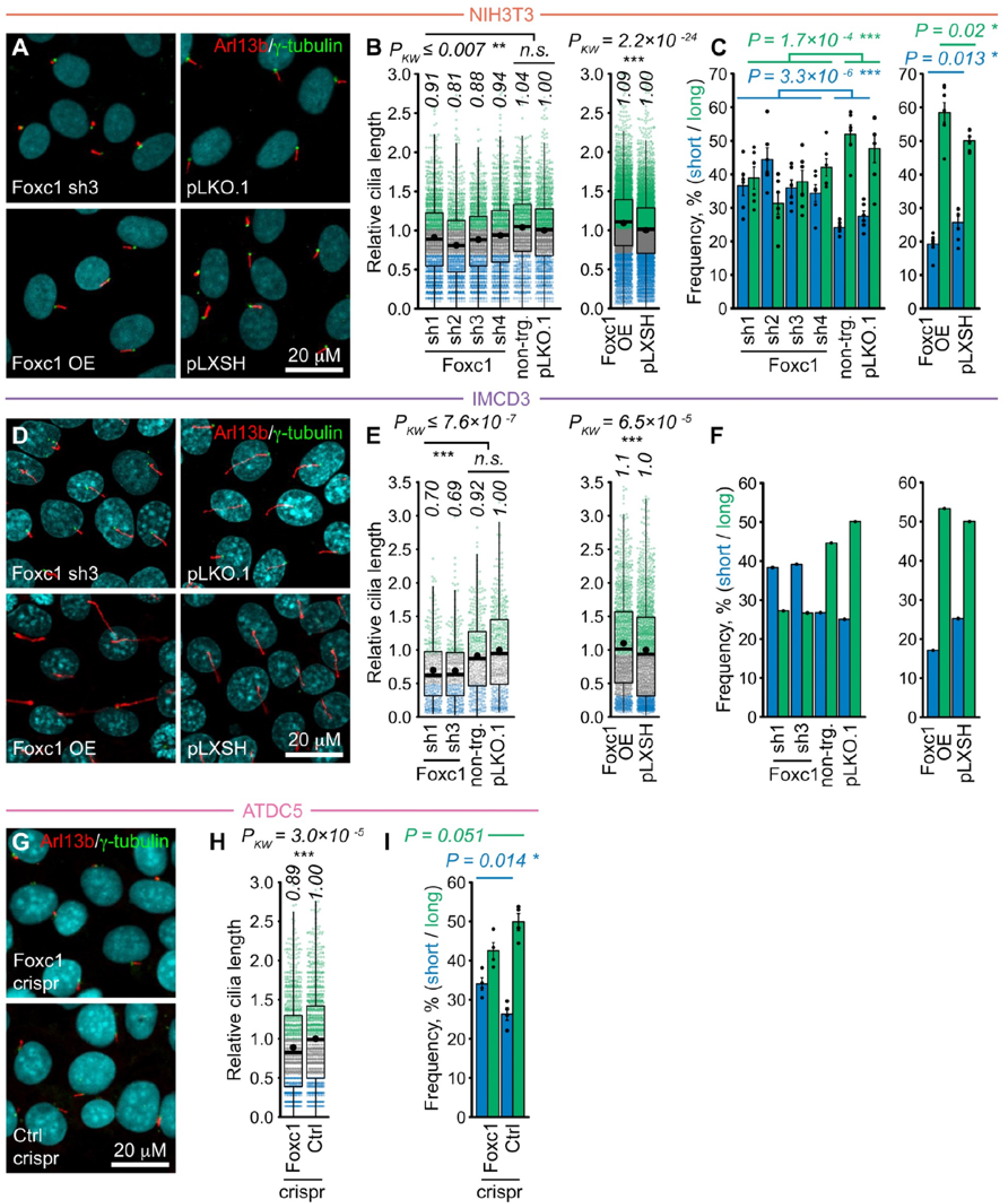
Altered levels of Foxc1 are associated with changes to cilia length in three cell types. (**A**) Representative images of Arl13b/*γ*-tubulin ciliary staining in NIH3T3 cells expressing a Foxc1-targeting shRNA, or with Foxc1 overexpression (OE) (vector controls: pLKO.1, pLXSH). (**B**) Quantification of cilia length as fold change relative to vector controls from n=6 (Foxc1 shRNA) or n=7 (Foxc1 OE) independent experiments. (**C**) Prevalence of short (length ≤ lower quartile of controls) and long (≥ median of controls) cilia in cells by condition. (**D**-**F**) In IMCD3 cells, Foxc1 knock-down and OE respectively reduce and increase cilia length; graphs depict individual length measurements and prevalence of short and long cilia, by condition. (**G**-**I**) In ATDC5 cells, the decreased cilia length induced by CRISPR mutagenesis of Foxc1 is attributable to an increased proportion of cells with short cilia. [Boxplots: median (black lines) and mean (black dots) values; statistical analyses: Dunn’s (*post hoc* Kruskal-Wallis) test. Barplots: mean values ± SEM; statistical analyses: two-way ANOVA (C shRNA), one-way ANOVA (C Foxc1 OE, I)].

### Altered Foxc1 dosage impacts cilia function, as illustrated by cilia-mediated Hh signaling

To test whether the alterations in axonemal length were associated with perturbed cilia-dependent signalling, we first assayed the activity of the Hedgehog pathway, that is mediated by the primary cilium in most vertebrate tissues (57–59). The Gli proteins (Gli1-Gli3) represent key effectors of the pathway, and the level of Gli1 was used as the initial readout. Under unstimulated conditions, cells overexpressing Foxc1 exhibit increased levels of Gli1 (Figure 3A). Conversely, cells with shRNA-mediated knockdown of Foxc1 accumulate lower levels of Gli1 on stimulation of Hh signaling with *Smoothened* agonist (SAG; Figure 3B-C). Alterations were not confined to Gli1, since significantly increased expression of *Gli2* (the main activator of mammalian Hh signalling) and *Ptch1* were observed in Foxc1-overexpressing fibroblasts (Figure 3D). To validate these findings, we next assayed the effect of manipulating Foxc1 levels using a second cell line: immortalized embryonic primary chondrocytes, that express high levels of endogenous Foxc1 (Supplemental Figure 2). In these mesenchymal-derived cells, shRNA-mediated knockdown of Foxc1 significantly decreased Gli1 expression (Figure 3E). Taken together, these data demonstrate that alterations to the level of Foxc1 perturb the *in vitro* expression of major Hh pathway components.

**Figure 3.**
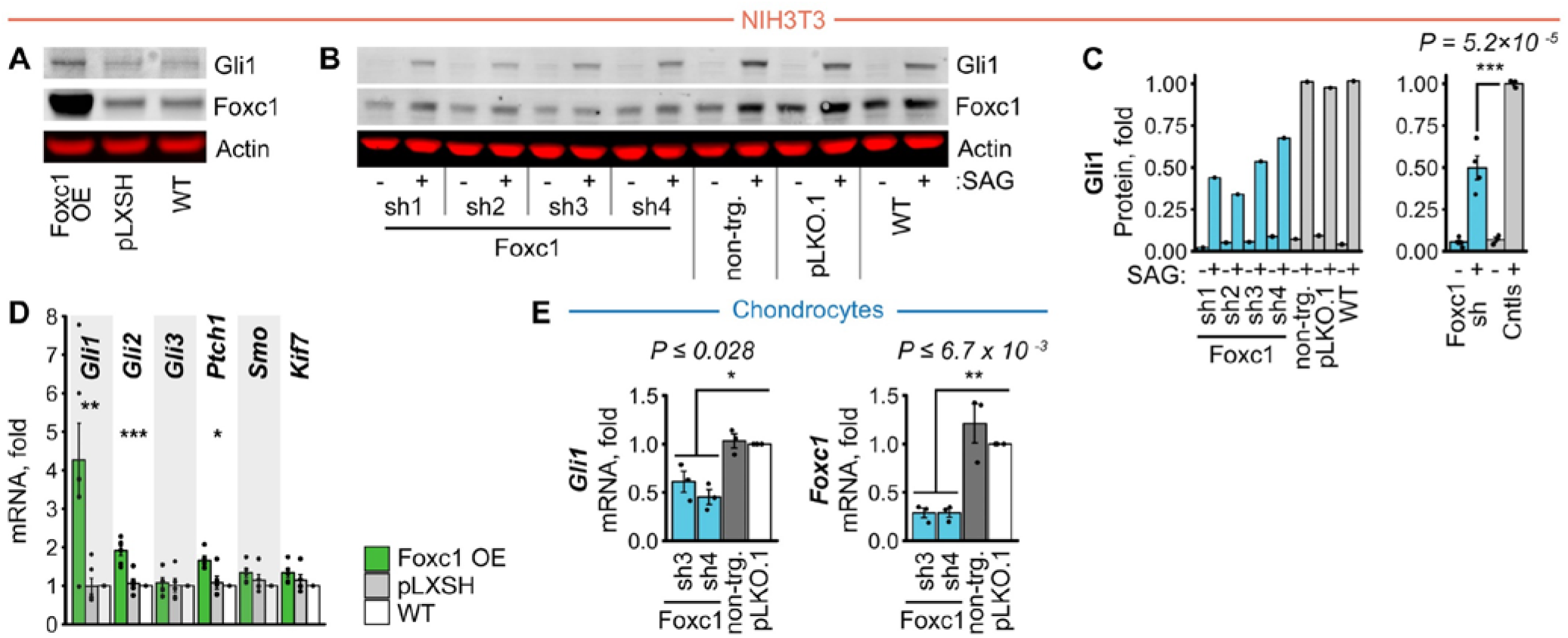
Altered levels of Foxc1 induce aberrant ciliary Hedgehog signaling. (**A-B**) In NIH3T3 cells, quantitative Western Immunoblots demonstrate altered levels of Gli1 protein with increased and decreased expression of Foxc1. Overexpression increases the basal level of Gli1 expression, while Foxc1 shRNA inhibition has a converse effect. (**C**) Overall, Foxc1 shRNA inhibition induces a 2-fold decrease in the level of Gli1 protein [Hh signaling stimulated in serum-starved cells with *Smoothened* agonist (SAG)]. (**D**) Cells overexpressing Foxc1 exhibit increased basal levels of *Gli1*, *Gli2* and *Ptch1* mRNA. (**E**) In immortalised E16.5 chondrocytes, that express high levels of Foxc1, shRNA inhibition of Foxc1 also decreased basal level of *Gli1* mRNA. [Barplots: mean values ± SEM; statistical analyses: Tukey HSD test *post hoc* one-way ANOVA].

### Foxc1 induces accumulation of Gli2 at the ciliary tip

A notable feature of vertebrate Hh signaling is that components change subcellular localization in response to ligand activation; as illustrated by the accumulation of Gli2 at the axonemal tip (60). Since this is essential to signal transduction, we established a clonal NIH3T3 cell line stably expressing Gli2-mGFP, and first demonstrated that it recapitulated the SAG-dependent accumulation of Gli1 protein observed in NIH3T3 cells (Figure 4B-D). The elevated basal level of Gli1 protein observed in these Gli2-mGFP cells, supports a cooperative effect of Foxc1 and Gli2 proteins on Hh pathway output (Figure 4A-D). Overexpression of Foxc1, significantly increased Gli2-mGFP accumulation at the ciliary tip, an effect that was further elevated by stimulation with SAG (~2-fold increase in mGFP signal intensity; *P*_*KW*_ = 6.2 × 10^−47^; Figure 4E-F). This enhanced axonemal tip accumulation of Gli2 is consistent with Foxc1 impacting a core cilia-mediated signalling pathway.

**Figure 4.**
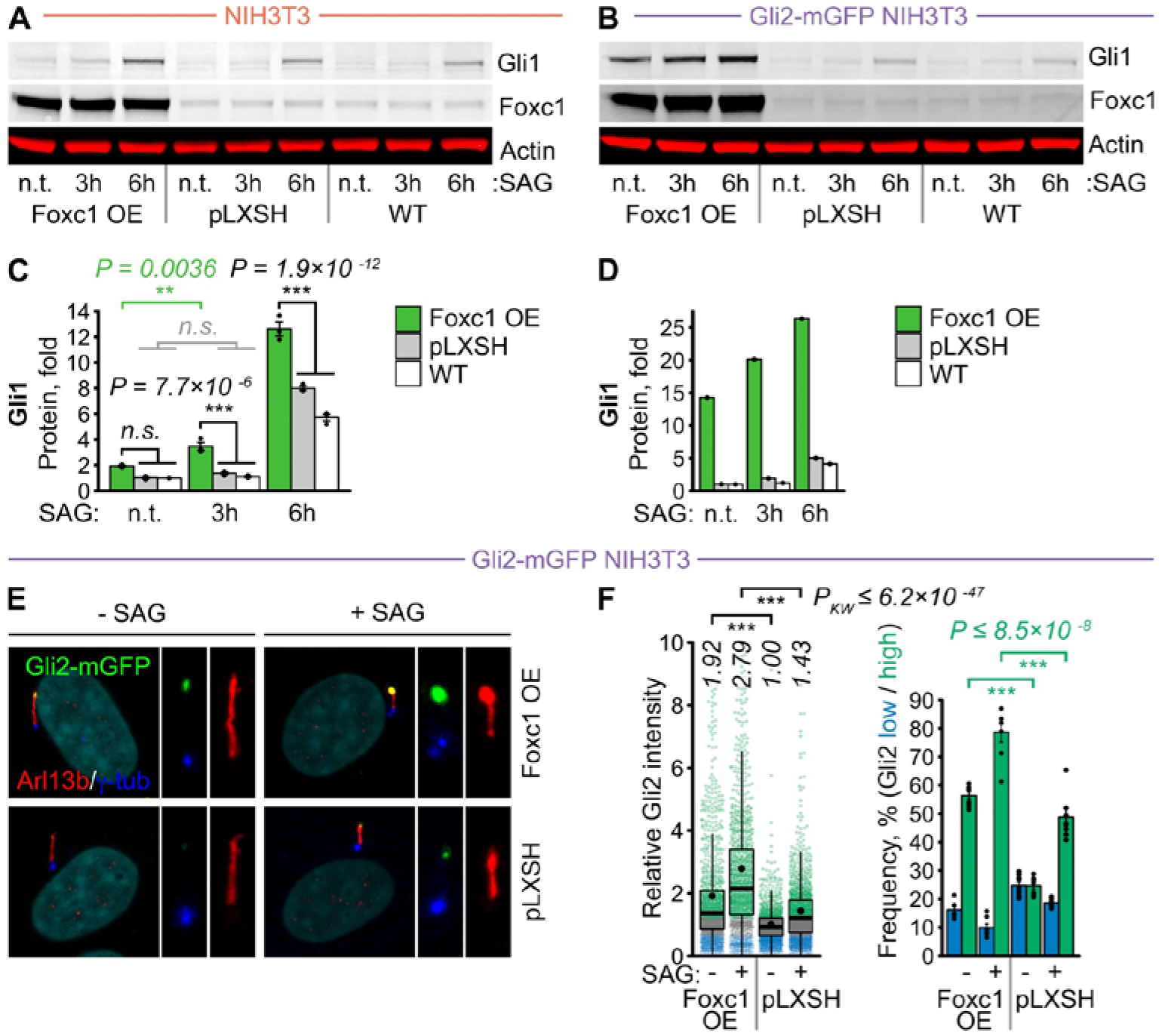
Foxc1 alters the dynamics of Hh signaling and enhances Gli2 accumulation at the ciliary tip. (**A-B**) Quantitative immunoblots demonstrate that overexpressing Foxc1 induces faster accumulation of Gli1 in serum-starved NIH3T3, and Gli2-mGFP NIH3T3 cells, subjected to stimulation with SAG. (**C**) Quantification of panel ‘A’, reveals a significant difference in Gli1 protein levels for Foxc1 OE condition at 3 and 6 hour time points. (**D**) In Gli2-mGFP NIH3T3 cells, quantification reveals an approximately two-fold higher level of Gli1 protein compared with Foxc1 OE NIH3T3 cells (>25-fold vs ~ 12-fold respectively, relative to non-treated WT controls). (**E**) Images show increased Gli2-mGFP accumulation at axonemal tips in cells overexpressing Foxc1, both before and after stimulation with SAG. (**F**) Quantification of ciliary tip Gli2-mGFP signal intensity expressed as fold change relative to mean untreated control [pLXSH]; together with frequencies of “bright” or “dim” (≥ upper quartile, ≤ lower quartile of control) axonemal tip Gli2-mGFP signal by condition from n=7 independent experiments. [Boxplots: median (black lines) and mean (black dots) values; statistical analysis: Dunn’s (*post hoc* Kruskal-Wallis) test. Barplots: mean values ± SEM; statistical analyses: Tukey HSD *post hoc* two-way (C) or one-way (F) ANOVA].

### Altered levels of Foxc1 impact platelet-derived growth factor signaling

Multiple developmental pathways, including WNT, Notch and several receptor tyrosine kinases, are partially mediated by cilia. Accordingly, we next assayed the PDGFR_α_ pathway to determine whether Foxc1’s effects on cilia-dependent signalling extend beyond Hedgehog signal transduction. Initiated at primary cilia under conditions of serum starvation, ligand stimulation by Platelet-derived growth factors (PDGF-A to C) induces phosphorylation of specific PDGFR_α_ tyrosine residues, such as pY754, that provide a readout of receptor activity. We observed that serum-starved NIH3T3 cells accumulate lower levels of PDGFR_α_ protein with either knockdown or overexpression of Foxc1 compared to controls; and when stimulated with cognate PDGF-AA ligand, increased and decreased levels of Foxc1 lead to reduced kinase activity of the PDGFR_α_ receptor. Foxc1 shRNA knockdown induced a two-fold reduction in levels of pY754 autophosphorylated PDGFR_α_ (Figure 5A-C), while Foxc1 overexpression resulted in a 1.5-fold reduction in total levels of PDGFR_α_ (*P* = ~10^−10^; Figure 5D-F). These data demonstrate that Foxc1 moderately affects protein levels of total PDGFR_α,_ while strongly impacting its ligand-dependent phosphorylation. These results accord with our assays of Hh signaling and demonstrate that *in vitro*, alteration to the level of Foxc1 affects multiple cilia-mediated signaling pathways.

**Figure 5.**
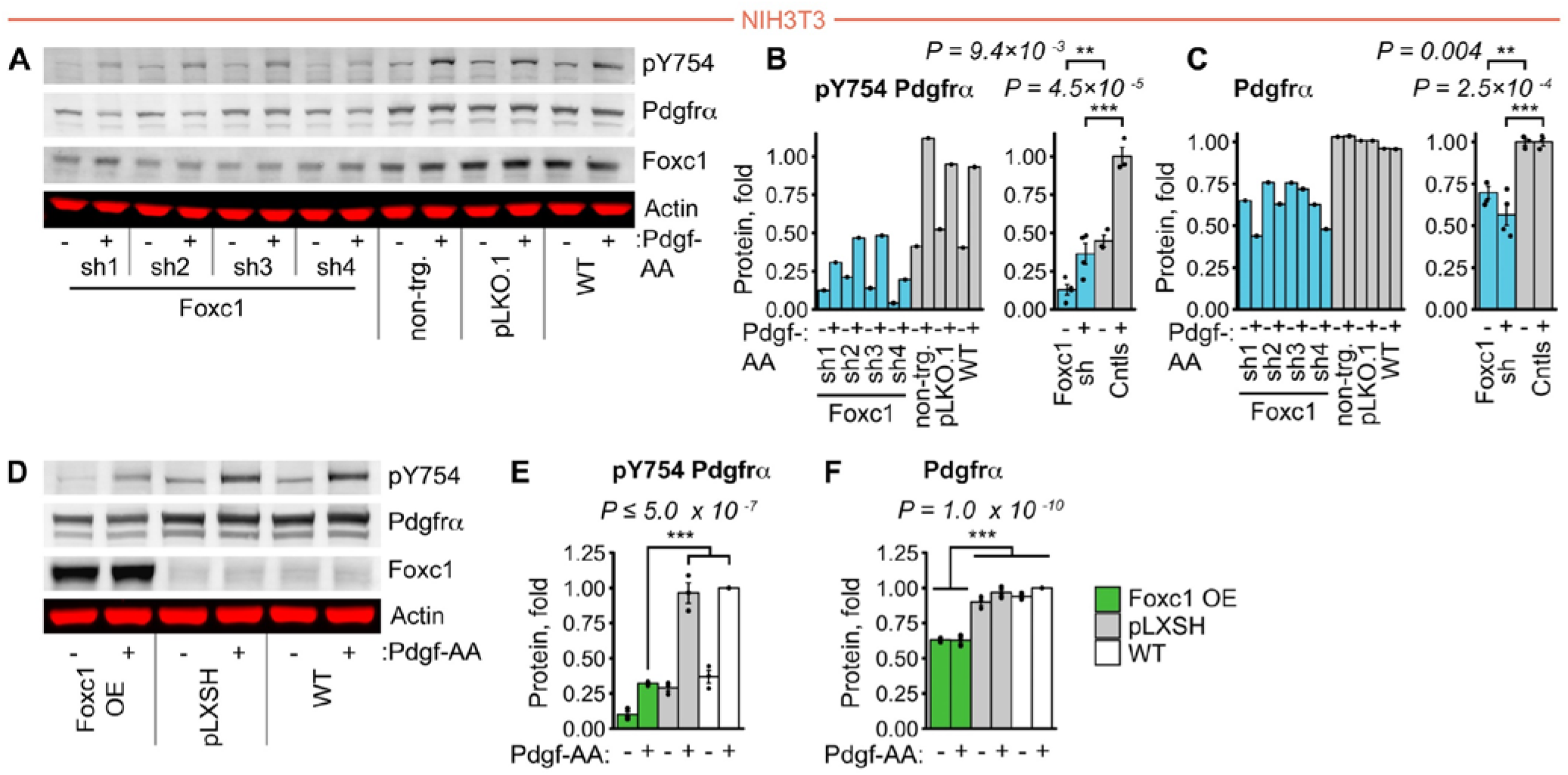
Altered levels of Foxc1 impact PDGFRα signaling. (**A**-**C**) Quantitative immunoblots demonstrate a consistent reduction in the level of auto-phosphorylated (pY754) and total PDGF Receptor alpha with Foxc1 shRNA inhibition (serum-starved NIH3T3 cells after 5 min stimulation with PDGF-AA ligand). (**D**-**F**) In cells overexpressing Foxc1, quantitative immunoblots reveal reduced levels of pY754 and total PDGFRα, indicative of impaired cilia-mediated PDGF signaling. [Barplots: mean values ± SEM; statistical analyses: Tukey HSD *post hoc* one-way (B,C,E) or two-way (F) ANOVA].

### Foxc1 mutation impacts choroid plexus structure and ciliation

To determine if comparable effects occur *in vivo*, we next investigated the CNS of Foxc1 murine mutants. We used homozygous embryos with a null *Foxc1*^*lacZ*^ deletion (hereafter *Foxc1*^−/−^), supplemented by embryos with complete *Cre*-mediated deletion of the *Foxc1* open reading frame (Foxc1^Δ/Δ^) (27, 61). At E14.5 and E15.5, *Foxc1*^−/−^ and Foxc1^Δ/Δ^ embryos display the expected severe dilation of the lateral ventricles together with hypoplasia of falx cerebri (Supplemental Figure 3A). In addition, we observed malformation of the choroid plexus in the lateral ventricles, evident by E15.5 as dysmorphic tortuous folding (Supplemental Figure 3B). This was associated with reduced size of the fourth ventricle, although the overall morphology of the fourth ventricle’s choroid plexus remained relatively preserved (Supplemental Figure 4).

Flow of cerebrospinal fluid (CSF) is essential to brain development and homeostasis (62), and perturbations are frequent causes of ventricular dilation and hydrocephalus. The choroid plexus, which is the primary source of CSF, comprises a multi-ciliated epithelial monolayer with a vascularized stromal core. Due to Foxc1’s causation of some forms of hydrocephalus, and changes to choroid plexus structure (Supplemental Figure 4), we examined ciliation in the lateral ventricles of E14.5 Foxc1^−/−^ mutants. In contrast to the unaltered cilia lining the ventricular ependyma, the length of the motile cilia lining the choroid plexus epithelium was prominently increased, demonstrated by immunostaining using antibodies to Arl13b, and separately acetylated tubulin (Figure 6A-B). Quantification revealed a 30% increase in axonemal length in *Foxc1*^−/−^ mutant embryos, primarily attributable to a greater prevalence of longer cilia (mean cilia length: *Foxc1*^−/−^ 2.08μm, Wildtype 1.59μm; *P* = 3.7 × 10^−4^; Figure 6C). Examination of ciliation at a later timepoint in E15.5 Foxc1^Δ/Δ^ embryos validated these findings (Figure 6D). Collectively, these data demonstrate that loss of Foxc1 impacts ciliation *in vivo*.

**Figure 6.**
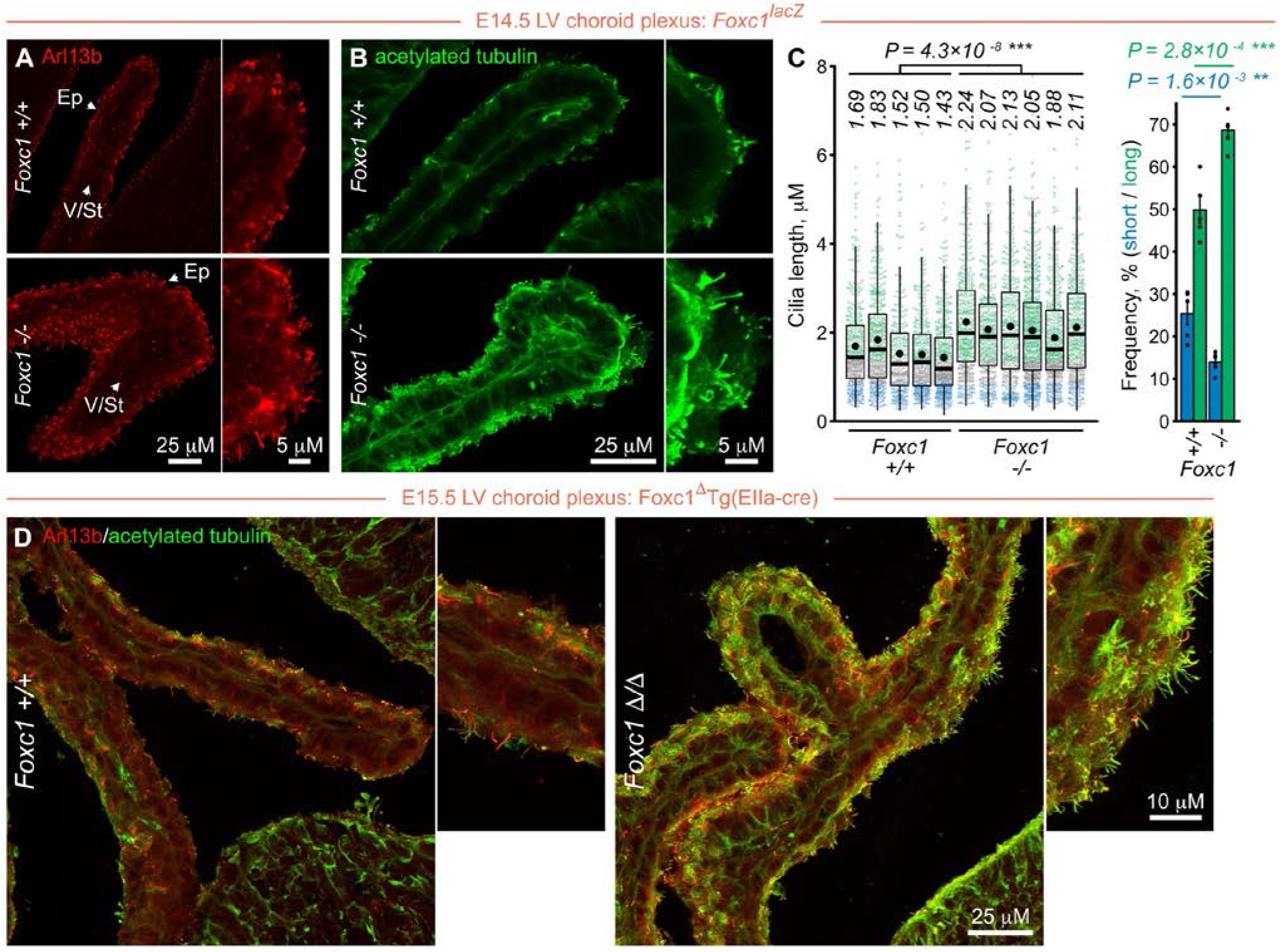
Altered ciliation in the choroid plexus of *Foxc1* mutant embryos. Upper panel (**A-D**): primary cilia length in choroid plexus epithelium of E14.5 *Foxc1*^−/−^embryos is increased relative to wild-type littermates: immunostaining performed to (**A**) Arl13b (ciliary membrane) and (**B**) acetylated tubulin (axonemal microtubules). (**C**) Quantification of individual cilia length for each embryo confirms a significant increase in the choroid plexus epithelium of *Foxc1*^−/−^mutants compared to wild-type controls, that is attributable to an increased proportion of long cilia in *Foxc1*^−/−^mutants. Lower panel (**E**) illustrates increased choroid plexus epithelium ciliation observed in E15.5 *Foxc1*^Δ/Δ^ embryos with *Cre-*induced deletion of *Foxc1*. Immunostaining performed to Arl13b and acetylated tubulin, in 2 *Foxc1* ^Δ/Δ^ and 2 *Foxc1*^+/+^ embryos. [Boxplot: median (black lines) and mean (black dots) values; statistical analysis: nested ANOVA. Barplot: mean values ± SEM; statistical analyses: one-way ANOVA. Ep, choroid plexus epithelium; V/St, choroid plexus vasculature / connective stroma].

### Foxc1 mutation leads to upregulation of Foxj1 expression in the choroid plexus

The increased axonemal length was unexpected, because the converse had been observed in cell culture. Potential explanations included the choroid plexus’ complexity compared to a single cell monolayer, and the possibility of differing effects on individual sub-populations of cilia. Accordingly, we assessed the level of Foxj1, which is a key regulator of multi-ciliation in the choroid plexus and other tissues. Immunohistochemistry on sections from wildtype E14.5 embryos, demonstrated the presence of Foxj1 protein in the nuclei of a small subset of choroid plexus epithelial cells. In contrast, in *Foxc1*^−/−^ mutants, the number of cells expressing Foxj1 and the level of Foxj1 expression, were both significantly increased, as illustrated by the 1.75-fold increase in the overall level of Foxj1 protein (*P* = 1.5 × 10^−3^, Figure 7A-C). Analysis of Foxj1 expression in the choroid plexus epithelium of E15.5 Foxc1^Δ/Δ^ embryos revealed similar changes (Figure 7D). Taken together, these data demonstrate that Foxc1 mutation impacts ciliation *in vivo*, and for a sustained interval, dysregulates expression of a paralog with essential roles in ciliogenesis. These effects likely reflect non-autonomous cell signaling, since *Foxc1* is expressed in the choroid plexus vasculature and stroma (Supplemental Figure 5) while the epithelium expresses *Foxj1*.

**Figure 7.**
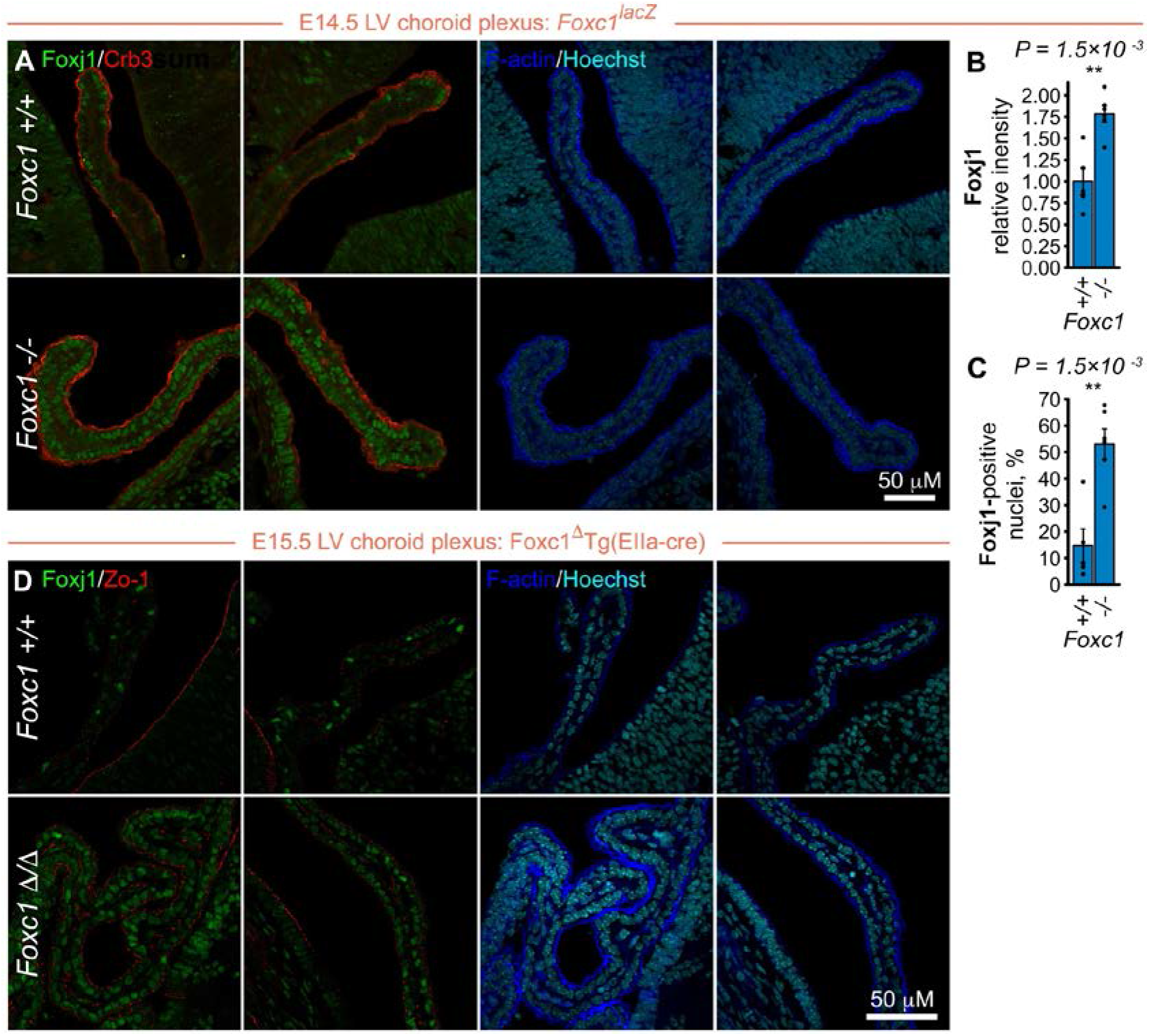
Increased expression of Foxj1 in the choroid plexus of *Foxc1* mutant embryos. Upper panel (**A-C**): Immunofluorescent analysis of E14.5 Foxc1^−/−^embryos demonstrates increased Foxj1 expression in the choroid plexus epithelium, and an increased proportion of Foxj1-positive nuclei. (**B**, **C**) There is a 1.75-fold increase in mean Foxj1 nuclear signal intensity, and the proportion of Foxj1-positive nuclei in the choroid plexus of Foxc1^−/−^embryos is significantly increased compared to wild-type controls. Lower panel (**D**) reveals an equal increase in Foxj1 expression in the choroid plexus of E15.5 *Foxc1*^Δ/Δ^ embryos after complete *Cre*-induced deletion of *Foxc1* [Barplots: mean values ± SEM; statistical analyses: one-way ANOVA].

## Discussion

This study presents evidence that in ARS, a neuro-developmental disorder characterized by heterogeneous systemic organ involvement (25, 43, 63–65), mutation of Foxc1 induces cilia dysfunction. Although *FOXC1* is responsible for a remarkable range of diseases, the mechanism by which mutation causes phenotypes as diverse as glaucoma, cancer and stroke, was undetermined. Our findings of a mechanistic connection with cilia offer an explanation for how alterations to a single gene induce seemingly unrelated phenotypes (pleiotropy), and derive from three lines of evidence. First, in fibroblast and renal cell lines, increased and decreased levels of Foxc1 induce reciprocal effects on axonemal length. These relatively small structural changes, resolved by developing automated quantification of large numbers of axonemes, were associated with more substantial (> two fold) alterations to cilia-mediated (Hh and PDGFRα) signaling. Such disproportionately severe functional consequences associated with relatively subtle structural changes, reiterate findings in Bardet-Biedl Syndrome, where some BBS mutants assemble visibly normal cilia but exhibit strong Hh-related patterning defects [reviewed in (66)], or loss of PGFFRα signaling (67). Second, our data demonstrate perturbed partitioning of proteins within the primary cilium, a process fundamental to cilia-mediated signaling. As illustrated by altered Gli2 accumulation at the tip of the axoneme, at a minimum the consequences of increased Foxc1 expression disrupt the tight regulation of Hh signaling, a developmental pathway with major additional roles in the maintenance and regeneration of adult tissues. The third line of evidence stems from alterations to cilia length in the murine choroid plexus.

Ciliation in the cerebral ventricles is dynamic, and surprisingly complex. At the E14.5 - E15.5 time points studied, the lateral ventricles are lined by neural progenitor cells with non-motile primary cilia: motile multi-ciliation develops later prenatally, extending into the early postnatal period (68, 69). In contrast, individual cells of the choroid plexus epithelium have multiple motile primary cilia (9+0 microtubule doublets), that exhibit a lower beat frequency and less motility than other multi-ciliated epithelia (70). Intriguingly, these cilia have variable axonemal architecture, with frequent ‘9+1’, and ‘9+2’ microtubules, in addition to the ‘9+0’ normally present in primary cilia (71). It was within this unique sub-population of cilia that altered cilia length was resolved in both *Foxc1*^−/−^ and Foxc1^Δ/Δ^ mutants, demonstrating that Foxc1 mutation perturbs ciliation *in vivo*. Since altered ventricular cilia number, structure or organization cause hydrocephalus (68, 72–74), these data offer a logical explanation for the ventricular dilation and occasional hydrocephalus observed in patients with heterozygous *FOXC1* mutation, and the congenital hydrocephalus apparent in homozygous murine and zebrafish mutants. Because choroid plexus cilia contribute to the control of CSF production, rather than producing measurable CSF flow (71), the changes observed are most consistent with compromised mechano- or chemosensation impairing regulation of CSF secretion.

Axonemal length represents a widely used, but indirect readout for ciliary function. Plausibly, the discordant readings of shortening in cultured single cell monolayers compared to lengthening *in vivo*, may reflect *Foxc1*’s regulation of multiple signalling pathways, including RA, PDGFRα, BMP and Wnt. Accordingly, we evaluated *Foxj1* to provide mechanistic insight for the *in vivo* changes. This *Forkhead* transcription factor has a fundamental role in motile ciliogenesis, and lies close to the apex of the genetic hierarchy that controls the development of multi-ciliation (11–13). In the lateral ventricles, *Foxj1* expression precedes outgrowth of monocilia and later ependymal cell multiciliation (75), and in the choroid plexus, like other tissues, controls ciliogenesis in multi-ciliated cells. Consequently, the increased protein expression of Foxj1 with *Foxc1*^−/−^ mutation links *Foxc1* with a key ciliary transcriptional regulator (11–13), so providing one potential mechanism for the axonemal length changes *in vivo*. The relationship between these two paralogs is of interest, due to numerous *Fox* genes having compensatory and overlapping developmental functions, while a subset exhibit antagonistic roles in the establishment of positional information (76–82). Our *in vitro* data support a *Foxc1*-dependent cell autonomous effect on cilia-mediated signaling, evident from the altered axonemal length and Gli2 partitioning in cells that do not express *Foxj1*, plus comparable changes in primary chondrocytes and cells with CRISPR-induced *Foxc1* mutation (Figures 2-4). Our *in vivo* findings are consistent with a model in which reduced expression of *Foxc1* within the vascular stroma of the choroid plexus, disrupts *Foxc1*-dependent signalling to the adjacent epithelium, impacting the normal expression of *Foxj1*. The relatively sustained duration of effect may facilitate future studies to identify mediators of the proposed non-cell autonomous signalling, that would enhance understanding of a tissue with fundamental roles regulating brain development and function. Equally, evidence that two *Fox* genes control aspects of choroid plexus ciliogenesis, supports the possibility that similar mechanisms are employed reiteratively in other tissues.

The data presented also provide a clinical explanation for the pleiotropic phenotypes associated with *FOXC1* mutation. Pleiotropy is an established feature of ciliopathies, and can result in a spectrum that extends from involvement of individual tissues to diverse combinations of organs (83). Consistently, the *Foxc1* murine mutant *congenital hydrocephalus* was originally reported as one of the earliest examples of a pleiotropic mutation variably altering a large number of organ systems (27, 40). This variable severity in patients and model organisms (27, 30, 38, 84–87), coupled with incremental identification of new phenotypes over more than a decade (29, 43, 88, 89), may explain the delay in discerning a ciliary contribution. The ARS phenotype most suggestive of a ciliary component was cerebellar hypoplasia (43), which reflects the reliance of cerebellar granule cell precursor proliferation on correct levels of Hh signaling (90, 91). Multiple other anomalies (Supplemental Table 1) are prevalent in the ciliopathy spectrum, consistent with the premise that ciliary dysfunction may contribute to such ARS phenotypes. Indeed, while this manuscript was being revised to include analyses of Foxc1^Δ/Δ^ embryos, demonstration that ablating cilia in the murine neural crest phenocopied the ocular mal-development characteristic of ARS (92), elegantly linked this phenotypic spectrum with altered ciliation. The clinical implications may extend beyond pediatric disease, to late onset disorders caused by *FOXC1*. For instance, Hh signaling’s maintenance of vascular integrity (93, 94), and pericyte recruitment (93–96), may plausibly explain *FOXC1*’s involvement in cerebral small vessel disease and stroke (25). Equally, dysregulated *FOXC1* expression is a poor prognostic factor in tumours such as breast cancer (97). The more invasive cell behaviours and metastasis that are associated with Foxc1’s aberrant activation of the Hedgehog pathway (98, 99) are thus rationally explicable in terms of ciliary dysfunction.

In conclusion, we have shown that alterations to the level of Foxc1 affect ciliation *in vitro* and *in vivo*, with evidence of both cell autonomous and non-cell autonomous mechanisms. These findings increase the number of *Forkhead* genes and clades with cilial functions, supporting sub-functionalization across a larger proportion of this transcription factor family. The impact on the expression of a master regulator of motile ciliogenesis in murine embryos with targeted and *cre*-mediated deletion of *Foxc1*, is particularly intriguing and signifies that in the choroid plexus a balance between Foxj1 and Foxc1 regulates properties of cilia. Such novel findings should encourage evaluation of the hypothesis that comparable transcriptional complexity is present in other multi-ciliated tissues. Overall, we propose that altered ciliation and cilia-mediated signaling contributes to the pleiotropic phenotypes induced by *FOXC1*.

## Methods

### Plasmids, antibodies and other reagents

Retroviral plasmid for stable expression of non-tagged mouse *Foxc1* ORF was created by InFusion HD® subcloning into pLXSH vector. mGFP-Gli2-pCEFL vector for expression of mouse Gli2 N-terminally tagged with monomeric GFP (mGFP) was a gift from Dr. Philip Beachy (Stanford University School of Medicine, Stanford, CA). Lentiviral shRNA plasmids from TRC1.5 mouse MISSION® shRNA libraries were obtained from the RNAi Screening Core (Li Ka Shing Institute of Virology, University of Alberta). Detailed information on all ORF and shRNA vectors, complete list of the antibodies, primers, as well as other reagents used in the study are provided in Supplemental methods (Tables S3 – S7).

### Cell lines

NIH3T3 mouse fibroblasts and derivative cell lines were grown at in DMEM supplemented with 10 % FBS; mIMCD3 inner medullary collecting duct cells – in DMEM:F12 supplemented with 10 % FBS; ATDC5 chondrogenic cells and derivative lines – in DMEM:F12 supplemented with 5% FBS and 2 mM GluaMAX-I. All cell lines were kept at 37°C in 5% CO_2_, in growth media containing 100 U/ml penicillin and 100 μg/ml streptomycin. Clonal NIH3T3 cell line stably expressing Gli2-mGFP was generated by transfection of parental cells with subsequent selection in 800 ug/ml Genetecin. NIH3T3 and Gli2-mGFP NIH3T3 cells stably expressing *Foxc1* were made by transduction with *Foxc1* ORF retroviral particles followed by selection in 100-500 ug/ml Hygromycin. Selection of NIH3T3 and mIMCD3 cells transduced with lentiviral shRNA particles was performed in 2.5 ug/ml Puromycin. Clonal ATDC5 cell line with mutation to *Foxc1* (*Foxc1*^*c.345_355del/c.353_356del*^) was generated using Alt-R Crispr-Cas9 system (IDT) (see Supplemental methods for details).

### Measurement of cilia length

Measurement of cilia length in NIH3T3 cells was performed after 20 h starvation in DMEM without FBS. In brief, after fixation in Dent’s solution, cilia and basal bodies of cells were immunofluorescenty labelled using anti-Arl13b and anti-*γ*-tubulin antibodies respectively (see Supplemental methods for details). Images were collected using Zeiss LSM 700 laser scanning confocal microscope and subjected to quantification of cilia length with automatic Cell Profiler (100) (the Broad Institute, Cambridge, MA) software-based image analysis pipeline. Measurements of cilia length in ATDC5 (automatic, Cell Profiler) and mIMCD3 (manual, using Fiji software (101)) were performed under conditions of growth in complete media with 10% FBS. For analyses of cilia length, short cilia were defined as having a length ≤ lower quartile of controls, and long cilia as ≥ median of controls.

### qPCR analyses

Total RNA was isolated with RNeasy Plus Mini Kit (Qiagen), quantified and used for cDNA synthesis with Primescript RT Master Mix (Clontech). qPCR reactions were run with SYBR® Premix Ex Taq (Tli RNAse H Plus) master mix (Clontech) on LightCycler® 96 Instrument and analysed using LightCycler® 96 Application (Roche Life Science). Primer sets used are provided in Supplemental methods (Table S7).

### Quantitative western blotting

Cells were lysed in 1.5% SDS lysis buffer (50 mM Tris pH 7.5, 150 mM NaCl, 1 mM EDTA, 1.5% SDS) supplemented with protease/phosphatase inhibitor cocktail (1 mM PMSF, 0.5 mM Na_3_VO_4_, 5 mM NaF, 10 mM β-glycerophosphate, 10 μg/ml aprotinin and 10 μg/ml leupeptin) and passed through QIAshredder columns (Qiagen). Obtained protein samples were normalised, resolved by SDS-PAGE, transferred to Immobilon-FL PVDF membranes (EMD Millipore) and blocked with Odyssey® TBS Blocking Buffer (Li-Cor). Next membranes were incubated with relevant primary antibodies, followed by IRDye-conjugated secondary antibodies. Resulting membranes were scanned with Odyssey® Imaging System (Li-Cor). List of primary and secondary antibodies used can be found in Supplemental methods (Table S5).

### Measurement of ciliary Gli2 accumulation

Accumulation of Gli2 at the cilia tips was measured in NIH3T3 Gli2-mGFP cells following 20 h starvation in DMEM medium without FBS, and subsequent stimulation with *Smoothened* agonist (SAG). After fixation in Dent’s solution, cells were immunofluorescenty labelled using anti-Arl13b and anti-*γ*-tubulin antibodies respectively (see Supplemental methods for details). Images were collected using Zeiss LSM 700 laser scanning confocal microscope and quantified with automatic Cell Profiler (99) software-based image analysis pipeline.

### Experimental Animals

The Foxc1^−/−^ embryos carrying the null *Foxc1*^*lacZ*^ mutation were generated and genotyped as previously described (27, 84). Embryonic age was determined by defining noon on the day of vaginal plug as E0.5. Genotyping of embryos was performed by PCR as described before (27) and additionally confirmed by qPCR end-point genotyping. Primers used for qPCR genotyping are provided in Supplemental methods (Table S7).

### Immunostaining of mouse tissues

Embryos were harvested at E14.5 in cold PBS and fixed in 4% paraformaldehyde (PFA) for 4 h at 4°C. Following fixation embryos were equilibrated in 20% sucrose in PBS, embedded in Clear Frozen Section Compound (VWR, Richmond, IL) and frozen in a dry ice bath. Coronal cryosections (12-16 μM) thick were made using Leica CM1900 cryostat and stored at - 85°C. Cilia in cornea and choroid plexus of the embryos were immunofluorescently labelled with anti-Arl13b and anti-acetylated tubulin antibodies. Cell polarity markers in cornea and meninges were labelled with anti-Prickle1 antibody and anti-Zo1-AF555 antibody conjugate. Nuclear Foxj1 and apical cell domain in choroid plexus were immunostained with anti-Foxj1 and anti-Crb3 antibodies respectively (see Supplemental methods for details). Images were collected using Zeiss LSM 700 laser scanning confocal microscope. Measurements of cilia length, cilia orientation and Prickle1 intensity in mouse tissues were performed using Fiji software (101). Measurements of nuclear Foxj1 intensity and quantification of Foxj1-positive nuclei were done with Fiji (101) and CellProfiler (100).

### Statistics

Statistical analyses were performed using RStudio version 0.99 software (RStudio Inc) running R language version 3.4.2 (The R Project for Statistical Computing). Analyses of statistical significance (P < 0.05) were performed with one-way ANOVA, two-way ANOVA, Tukey HSD test *post hoc* one-way and two-way ANOVA, nested ANOVA or Dunn’s (*post hoc* Kruskal-Wallis) test, as indicated for specific experiments. Bar plots show mean values ± SEM. Box-whisker plots show quartiles, median (black lines) and mean (black dots) values. Significance codes *** P < 0.001, ** P < 0.01, * P < 0.05.

### Study Approval

Ethical approval was provided by the University of Alberta Health Research Ethics Board, with written informed consent received from all participants prior to their inclusion in the study. Animal experiments were approved by the IACUC of the University of Alberta.

## Author Contributions

OJL and SH conceived the study. SH performed and analyzed the majority of the experiments. PC, SVB, CRF and FBB contributed to specific experiments and data analysis. TK and AJW were involved in study design and supervision. IMM provided additional funding for the study, and with JRA and RCR contributed patient phenotypic data and reagents. SH and OJL co-wrote the manuscript.

## Acknowledgements

We are grateful to the patients who participated in this study. We thank Sudipto Roy (National University of Singapore), Michael Walter (University of Alberta), Peter Carlsson (University of Gothenburg), and Valerie Wallace (University of Toronto) for critically reviewing the manuscript. Funding was provided by the Canadian Institutes of Health Research (CIHR) (MOP-133658) and Womens and Children’s Health Research Institute (to OJL), Natural Sciences and Engineering Research Council (to AJW) and Alberta Innovates Health Solutions (to IMM).

---

**Supplemental Table 1.** Prevalence of phenotypes with *FOXC1* mutation and CNV. Cardiac anomalies included atrial septal defects, mitral and tricuspid incompetence and a bicuspid aortic valve; 19 of the 28 patients with glaucoma required surgery. The different denominators in the upper (n=41) and lower (n=26) portions of the table reflect the number of patients who had cerebral imaging performed (MRI, n=24; CT, n=2).

|  | *FOXC1*<br>mutation<br>(n=19) | Segmental<br>deletion<br>(n=5) | Segmental<br>duplication<br>(n=17) | Total<br>(n=41) |
| --- | --- | --- | --- | --- |
| Glaucoma | 13 | 2 | 12 | 28 |
| Dental Anomalies | 7 | 1 | 2 | 10 |
| Umbilical Hernia | 1 | 0 | 0 | 1 |
| Mid-facial Hypoplasia | 9 | 0 | 2 | 11 |
| Hypertelorism | 5 | 1 | 0 | 6 |
| Auditory Impairment | 7 | 1 | 2 | 10 |
| Cardiac Anomalies | 10 | 1 | 0 | 11 |
| Skeletal Anomalies | 5 | 1 | 1 | 7 |
| Polydactyly | 1 | 0 | 0 | 1 |
| Cerebral Small Vessel Disease | 12 | 5 | 8 | 25 of 26 |
| Cerebellar Hypoplasia | 1 | 3 | 0 | 4 of 26 |
| Hydrocephalus | 1 | 1 | 0 | 2 of 26 |
| Ventriculomegaly | 1 | 2 | 0 | 3 of 26 |

**Supplemental Figure 1.**
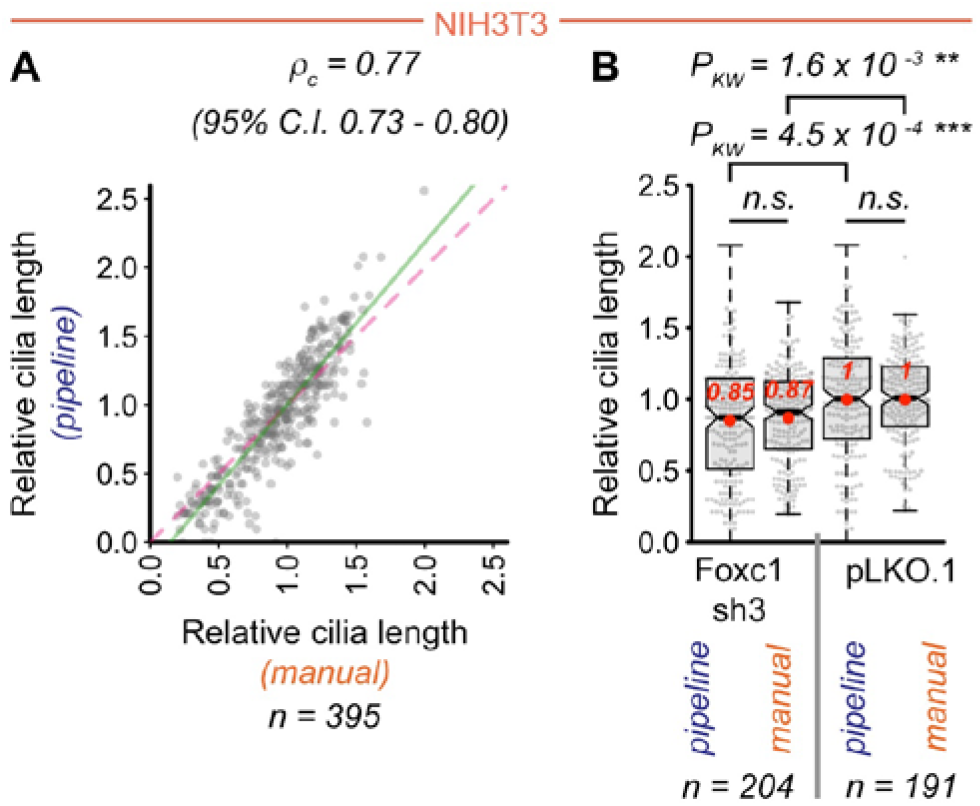
Quantification of cilia length in cells with altered Foxc1 dosage. (**A**) Concordance between automatic and manual quantification of cilia length in NIH3T3 cells. Individual axonemal length was quantified using Cell Profiler-based automatic pipeline and manually in 395 cells. Graph shows automatic pipeline (Y-axis) and corresponding manual (X-axis) measurements relative to vector control (pLKO.1) plotted against each other. Lin’s concordance correlation coefficient calculated for pipeline vs manual measurements (*ρ*_c_ = 0.77) shows moderate level of agreement between the two methods [line of perfect concordance (dashed pink); line of linear regression of automatic vs manual quantification values (solid green); Lin LIK, Hedayat A, and Wu W. *Statistical tools for measuring agreement.* New York: Springer; 2012]. (**B**) Relative cilia length (gray dots) in cells expressing a Foxc1-targeting shRNA (Foxc1 sh3) or vector control (pLKO.1) quantified using automatic pipeline and manually. Note: a) lack of significant difference between automatic vs manual quantification for both conditions, b) comparable statistically significant difference between Foxc1 shRNA and control conditions for both quantification methods.

**Supplemental Figure 2.**
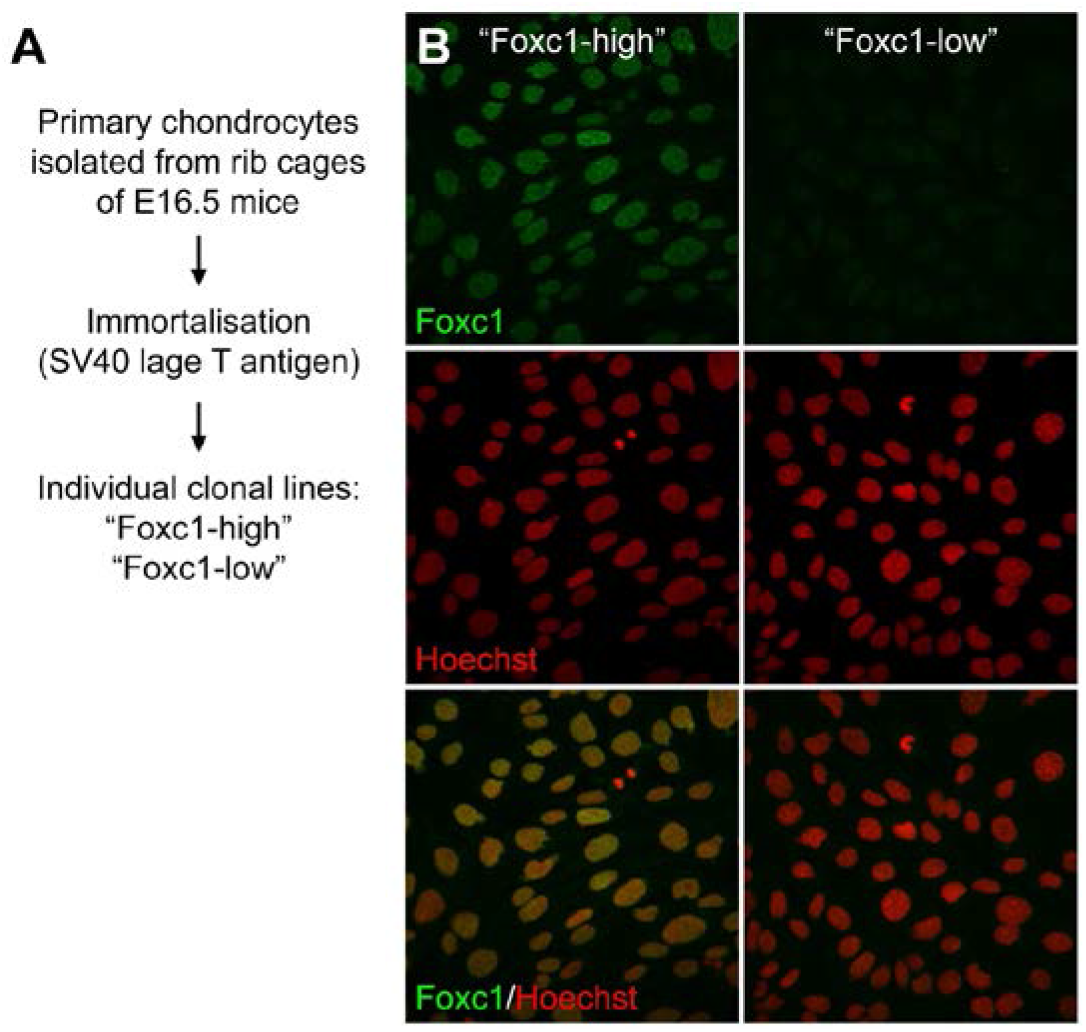
Foxc1 expression in immortalised mouse chondrocytes. (**A**) Primary chondrocytes were isolated from rib cages of E16.5 mice and immortalised by retroviral transduction with SV40 large T antigen. Individual clonal cell lines were obtained and assayed for Foxc1 expression. (**B**) Microscopic images exemplify difference in nuclear immunofluorescent staining of Foxc1 in immortalised chondrocyte lines expressing high and low levels of the protein respectively.

**Supplemental Figure 3.**
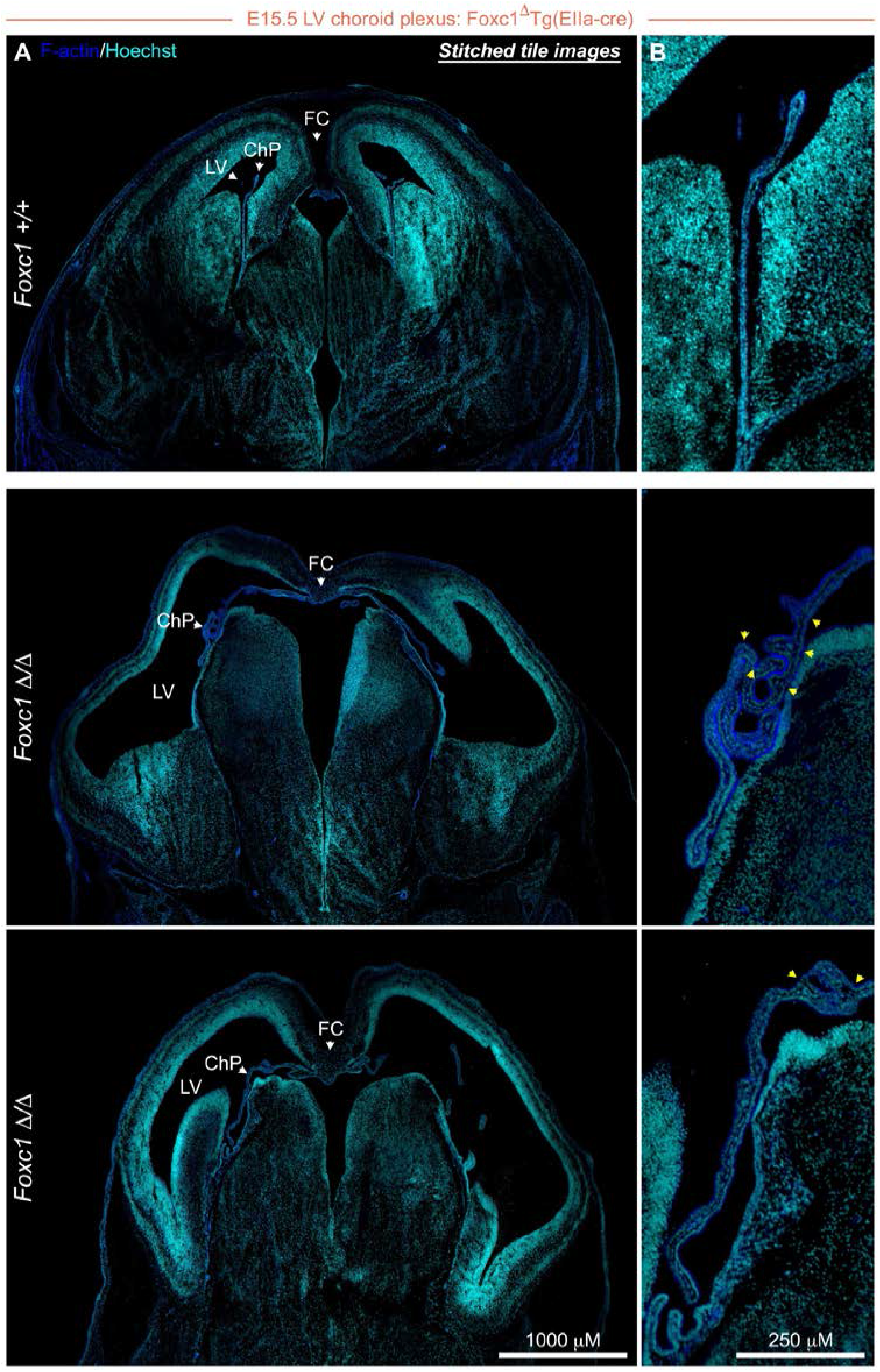
Morphology of lateral ventricles and choroid plexus in *Foxc1* null embryos. (**A**) Coronal sections illustrate the morphology of the lateral ventricles in E15.5 *Foxc1*^Δ/Δ^ embryos upon complete deletion of *Foxc1* ORF: note the dilation of the ventricular lumen and underdeveloped falx cerebri relative to wild type controls. (**B**) Insets highlight the substantial dysmorphism of the lateral ventricle choroid plexus, with extensive folding of choroid plexus tissue in *Foxc1*^Δ/Δ^ mice (yellow arrows). [LV, lateral ventricle; ChP, choroid plexus; FC, falx cerebri].

**Supplemental Figure 4.**
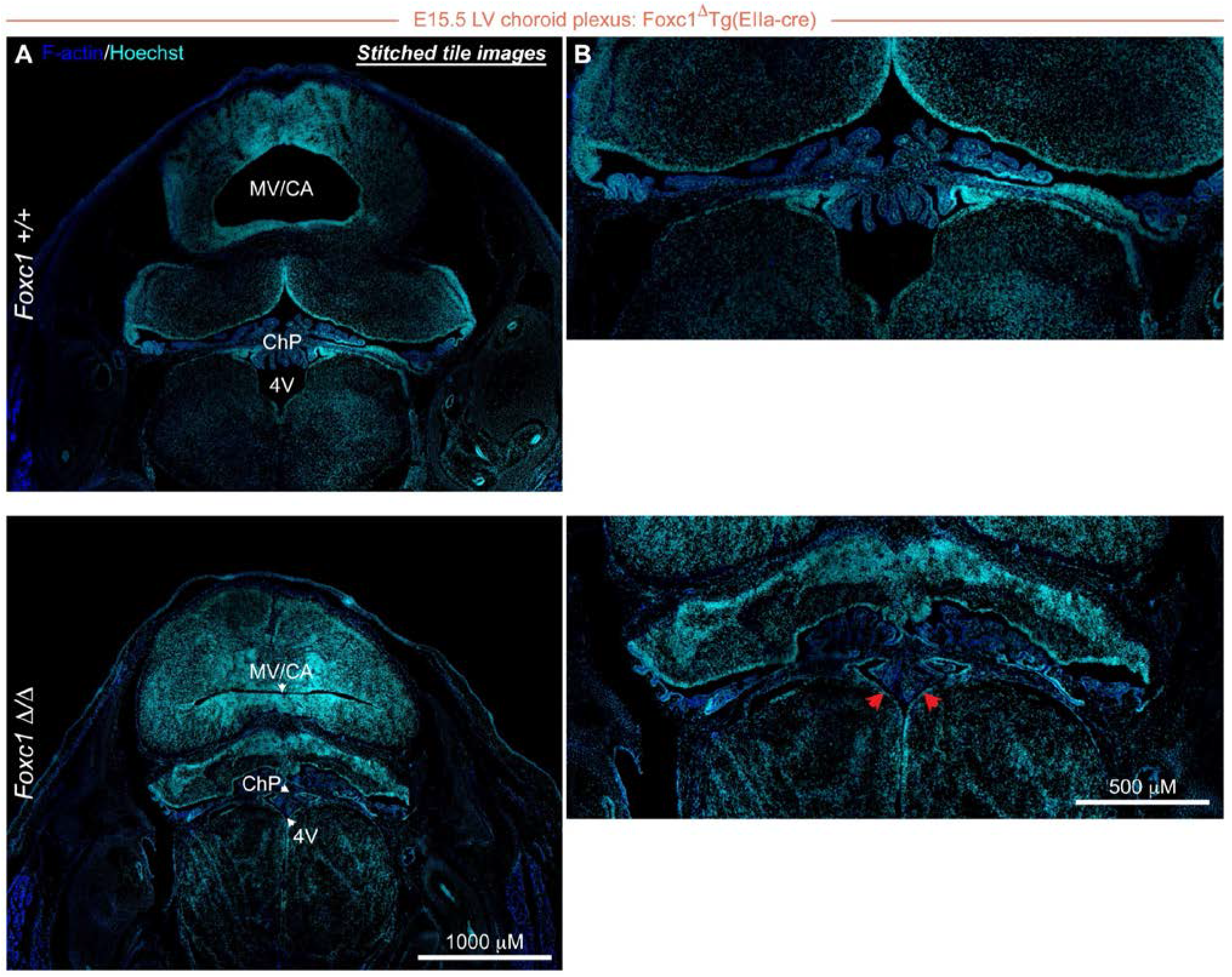
Morphology of the fourth ventricle and choroid plexus. (**A-B**) Coronal sections illustrate the altered size of the fourth ventricle, delineated by red arrows, with relatively preserved choroid plexus morphology at E15.5 in *Foxc1*^Δ/Δ^ embryos. [4V, fourth ventricle; ChP, choroid plexus; MV/CA, mesencephalic vesicle / cerebral aqueduct].

**Supplemental Figure 5.**
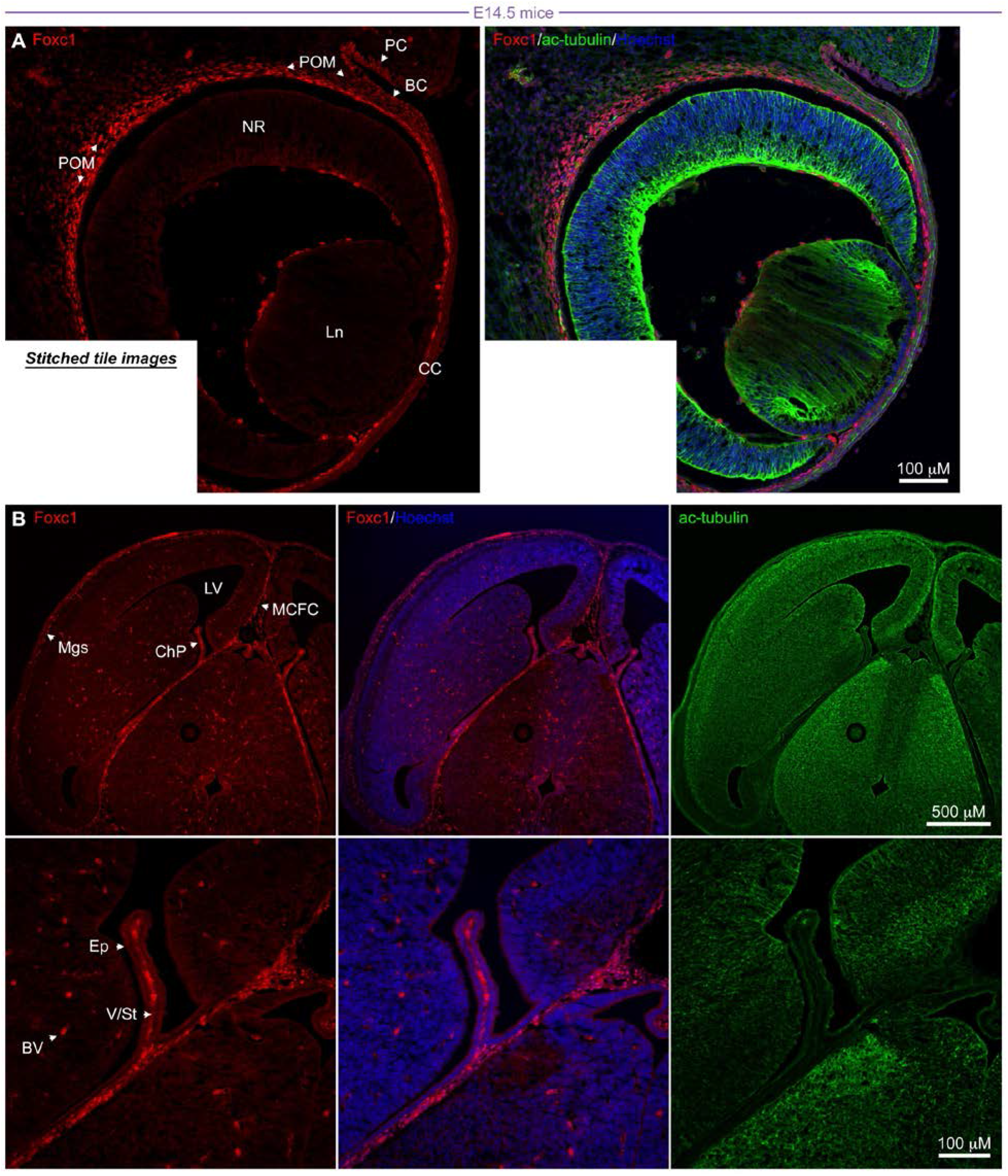
Expression of Foxc1 in the periocular mesenchyme and choroid plexus. (**A**) Immunofluorescent staining of ocular coronal sections of E14.5 murine embryos demonstrates high levels of Foxc1 protein in wild type periocular mesenchyme (POM). (**B**) Immunofluorescent analysis of horizontal sections through the lateral ventricles reveals high levels Foxc1 protein expression in the choroid plexus, brain vasculature and meninges E14.5. [Ln, lens; CC, central cornea; NR, neural retina; PC, palpebral conjunctiva; BC, bulbar conjunctiva; LV, lateral ventricle; ChP, choroid plexus; Ep, choroid plexus epithelium; V/St, choroid plexus vasculature / connective stroma; BV, brain vasculature; Mgs, meninges; MCFC, mesenchymal condensation forming falx cerebri].

